# Impact of urolithiasis and hydronephrosis on acute kidney injury in patients with urinary tract infection

**DOI:** 10.1101/2020.07.13.200337

**Authors:** Chih-Yen Hsiao, Tsung-Hsien Chen, Yi-Chien Lee, Ming-Cheng Wang

## Abstract

**Background:** Urolithiasis is a common cause of urinary tract obstruction and urinary tract infection (UTI). This study aimed to identify whether urolithiasis with or without hydronephrosis has an impact on acute kidney injury (AKI) in patients with UTI.

**Methods and findings:** This retrospective study enrolled hospitalized UTI patients who underwent imaging in an acute care setting from January 2006 to April 2019. Of the 1113 participants enrolled, 191 (17.2%) had urolithiasis and 76 (6.8%) had ureteral stone complicated with hydronephrosis. Multivariate logistic regression analysis showed that in UTI patients with urolithiasis, the presence of ureteral stone with concomitant hydronephrosis was an independent risk factor for AKI (odds ratio [OR] 2.345, 95% confidence interval [CI] 1.128–4.876, P = 0.023). In addition, urolithiasis was associated with an increased risk for AKI (OR 2.484, 95% CI 1.398–4.415, P = 0.002) in UTI patients without hydronephrosis.

**Conclusion:** The presence of ureteral stone with hydronephrosis increases the risk for AKI of UTI patients with urolithiasis, and urolithiasis remains a risk factor of AKI in UTI patients without hydronephrosis.

## Introduction

Urolithiasis is a widespread problem with a male preponderance, showing a lifetime prevalence of 10% in men and 5% in women [1, 2]. The worldwide prevalence of urolithiasis is also steadily increasing [1-3], partially owing to its strong association with metabolic syndrome conditions such as obesity, hypertension, and diabetes [4].

Urolithiasis can cause secondary complications, such as obstruction or urinary tract infection (UTI) [5]. Acute UTI may cause sudden deterioration of renal function, especially in the presence of urinary tract obstruction [6]. Our previous study showed that urolithiasis is a risk factor for acute kidney injury (AKI) among patients with UTI [7], and AKI has been associated with short- and long-term mortality [8]. Since AKI leads to poor outcomes, recognizing the risk factors for its development in patients with urolithiasis and UTI is crucial. However, no study has focused on whether urolithiasis without hydronephrosis is a risk factor for the development of AKI in patients with UTI, or whether hydronephrosis caused by ureteral stone is a risk factor for the development of AKI in UTI patients with urolithiasis. Hence, we conducted this study to investigate these important issues.

## Methods

### Patient selection

This retrospective observational study covered the period from January 2006 to April 2019 and was conducted at Chia-Yi Christian Hospital, a tertiary referral center in the southwestern part of Taiwan. This study was performed after obtaining ethical approval from the Institutional Review Board of Chiayi Christian Hospital (approval no. CYCH-IRB-2019061).

A total of 1113 consecutive hospitalized patients with UTI without any other concurrent infectious disease were enrolled. All enrolled patients fulfilled the following criteria: (a) with UTI symptoms and > 10^5^ colony-forming units/mL of bacterial isolates from a urine specimen; (b) had available antimicrobial susceptibility test results; and (c) underwent complete imaging examination, including ultrasonography, intravenous urography, or computed tomography, and had complete data of the required laboratory tests. Patients aged < 18 years, with concurrent infection other than UTI, without imaging studies, with incomplete data, or undergoing regular dialysis therapy were excluded from this study. The included patients were divided into three groups: UTI patients with no hydronephrosis or urolithiasis, UTI patients with ureteral stone and concomitant hydronephrosis, or UTI patients with urolithiasis but no hydronephrosis (Fig 1).

**Fig 1.**
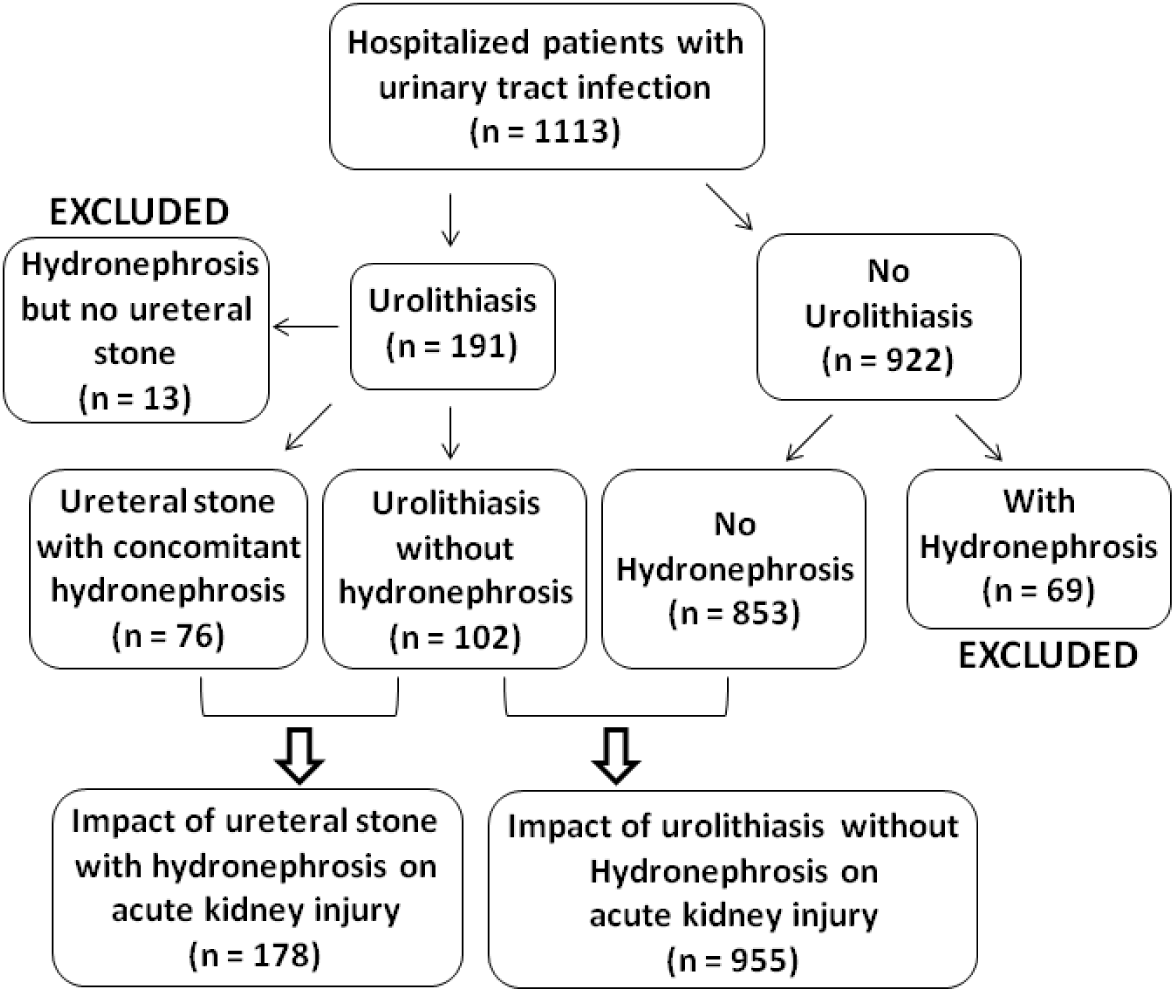
Flowchart of the selection of study subjects.

Inpatients were assessed using standard laboratory and diagnostic procedures. For further analysis, we collected clinical data in a standard form, including demographic characteristics (age and sex), comorbidities (coronary artery disease, congestive heart failure [CHF], stroke, diabetes mellitus [DM], or hypertension), presence of an indwelling Foley catheter, vital signs (blood pressure and ear temperature), laboratory results (white blood cell [WBC] count, platelet count, serum creatinine level, and estimated glomerular filtration rate [eGFR] at baseline and after hospitalization), prior history of UTI, existence of a urinary tract abnormality (urolithiasis or hydronephrosis), length of hospital stay, causative microorganisms (*Escherichia coli* [*E. coli*], *Proteus* species, *Klebsiella* species, *Enterococcus* species, and *Pseudomonas* species), and antimicrobial resistance pattern.

### Definition and examples of clinical covariates

The eGFR was determined using the Chronic Kidney Disease Epidemiology Collaboration creatinine equation and expressed in mL/min/1.73 m^2^ [9, 10]. AKI was defined as an increase in serum creatinine to ≥ 2.0 times the baseline value according to the Kidney Disease: Improving Global Outcomes Clinical Practice Guideline criteria for serum creatinine value for AKI stages 2 and 3 [11]. Hydronephrosis was defined as dilatation of the renal pelvis and calyces in imaging studies. Uroseptic shock was defined as sepsis-induced hypotension (systolic blood pressure [SBP] < 90 mmHg or mean arterial pressure < 70 mmHg or a decrease in SBP by > 40 mmHg lasting for at least 1 hour, despite adequate fluid resuscitation and in the absence of other causes of hypotension) [12]. An afebrile status was defined as a single temperature not increasing to > 38.3°C (101°F) [13]. Multiple drug resistance (MDR) was defined as non-susceptibility to at least three antimicrobial categories [14].

### Statistical analysis

All analyses were performed using SPSS version 17.0 (SPSS Inc., Chicago, IL, USA). Continuous variables are expressed as mean ± standard deviation, and categorical variables are expressed as number (percentage). Univariate analyses were performed using Student’s t-test for continuous variables, and the chi-square test and Fisher’s exact test for categorical variables. Multivariate logistic regression analyses were applied to identify risk factors associated with AKI during hospitalization. Only variables significant at the 0.15 level in univariate analysis were selected for the subsequent multivariate analysis. The goodness-of-fit of the logistic regression model was assessed using the Cox & Snell test, and the explanatory power was reported with Nagelkerke’s pseudo-*R*-square. P < 0.05 was considered statistically significant.

## Results

### Characteristics of the study population

The demographic and clinical characteristics of the 1113 hospitalized UTI patients included in this study are shown in Table 1. The mean age on admission was 66 ± 17 years. Most of the patients (816 [73.3%]) were women, and 375 (33.7%) had a history of prior UTI. Of the patients, 220 (19.8%) had uroseptic shock and 165 (14.8%) had AKI during hospitalization. The overall mortality rate was 0.6% (7/1113). Among UTI patients, 191 (17.2%) had urolithiasis and 76 (6.8%) had ureteral stone with hydronephrosis. Fig 2 shows a reduction in eGFR in different groups of hospitalized patients with UTI. The reduction in eGFR in UTI patients without urolithiasis or hydronephrosis, in those with urolithiasis but without hydronephrosis, and in those with ureteral stone and concomitant hydronephrosis was 13.45 (95% confidence interval [CI] 12.44–14.45), 20.54 (95% CI 16.34–24.73), and 27.65 (95% CI 21.77–33.53) mL/min/1.73 m^2^, respectively. Thirteen urolithiasis patients who had hydronephrosis but with no ureteral stone identified in imaging studies and 69 patients who did not have urolithiasis but had hydronephrosis were excluded from further analysis (Fig 1).

**Table 1.** Characteristics of hospitalized patients with urinary tract infection with respect to urolithiasis

|  | All<br>(n = 1,113) | Non-Urolithiasis<br>(n=922) | Urolithiasis<br>(n=191) | P-value |
| --- | --- | --- | --- | --- |
| Age (years) | 66±17 | 66±18 | 67±14 | 0.413* |
| Gender (male) | 297 (26.7) | 231 (25.1) | 66 (34.6) | 0.007<sup>¥</sup> |
| Diabetes mellitus | 478 (42.9) | 391 (42.4) | 87 (45.5) | 0.425<sup>¥</sup> |
| Hypertension | 584 (52.5) | 476 (51.6) | 108 (56.5) | 0.215<sup>¥</sup> |
| Congestive heart failure | 47 (4.2) | 38 (4.1) | 9 (4.7) | 0.712<sup>¥</sup> |
| Coronary artery disease | 122 (11.0) | 100 (10.8) | 22 (11.5) | 0.788<sup>¥</sup> |
| Stroke | 237 (21.3) | 198 (21.5) | 39 (20.4) | 0.746<sup>¥</sup> |
| Prior history of UTI |  |  |  | 0.091<sup>¥</sup> |
| None | 738 (66.3) | 622 (67.5) | 116 (60.7) |  |
| Once | 212 (19.0) | 174 (18.9) | 38 (19.9) |  |
| Twice | 89 (8.0) | 72 (7.8) | 17 (8.9) |  |
| Thrice or more | 74 (6.6) | 54 (5.9) | 20 (10.5) |  |
| Indwelling Foley catheter | 70 (6.3) | 56 (6.1) | 14 (7.3) | 0.515<sup>¥</sup> |
| Afebrile | 405 (36.4) | 342 (37.1) | 63 (33.0) | 0.283<sup>¥</sup> |
| Bacteremia | 519 (46.6) | 408 (44.3) | 111 (58.1) | <0.001<sup>¥</sup> |
| Uroseptic shock | 220 (19.8) | 163 (17.7) | 57 (29.8) | <0.001<sup>¥</sup> |
| Acute kidney injury | 165 (14.8) | 105 (11.4) | 60 (31.4) | <0.001<sup>¥</sup> |
| Length of hospital stay (days) | 9±5 | 9±5 | 11±5 | <0.001* |
| All-cause in-hospital mortality | 7 (0.6) | 6 (0.7) | 1 (0.5) | 0.656<sup>¥</sup> |
| Hospitalized serum creatinine (mg/dL) | 1.59±1.60 | 1.52±1.52 | 1.93±1.88 | <0.001* |
| Baseline eGFR (mL/min/1.73 m<sup>2</sup>) | 76.12±30.27 | 76.57±30.80 | 73.97±27.58 | 0.281* |
| Mean white blood cell (10<sup>3</sup>/μL) | 13.31±6.22 | 13.21±6.03 | 13.82±7.07 | 0.212* |
| Platelets (10<sup>3</sup>/μL) | 204.64±111.06 | 206.99±116.05 | 193.3±82.16 | 0.121* |
| *Escherichia coli* | 876 (78.7) | 747 (81.0) | 129 (67.5) | <0.001<sup>¥</sup> |
| *Proteus* species | 34 (3.1) | 16 (1.7) | 18 (9.4) | <0.001<sup>¥</sup> |
| *Klebsiella* species | 78 (7.0) | 63 (6.8) | 15 (7.9) | 0.615<sup>¥</sup> |
| *Enterococcus* species | 39 (3.5) | 32 (3.5) | 7 (3.7) | 0.894<sup>¥</sup> |
| *Pseudomonas* species | 55 (4.9) | 45 (4.9) | 10 (5.2) | 0.837<sup>¥</sup> |
| Multidrug-resistance bacteria isolate | 390 (35.0) | 318 (34.5) | 72 (37.7) | 0.398<sup>¥</sup> |

\*: Student T-test or Mann-Whitney U-test; <sup>¥</sup>:Chi-Square test or Fisher's exact test UTI: urinary tract infection; eGFR: estimated glomerular filtration rate.

**Fig 2.**
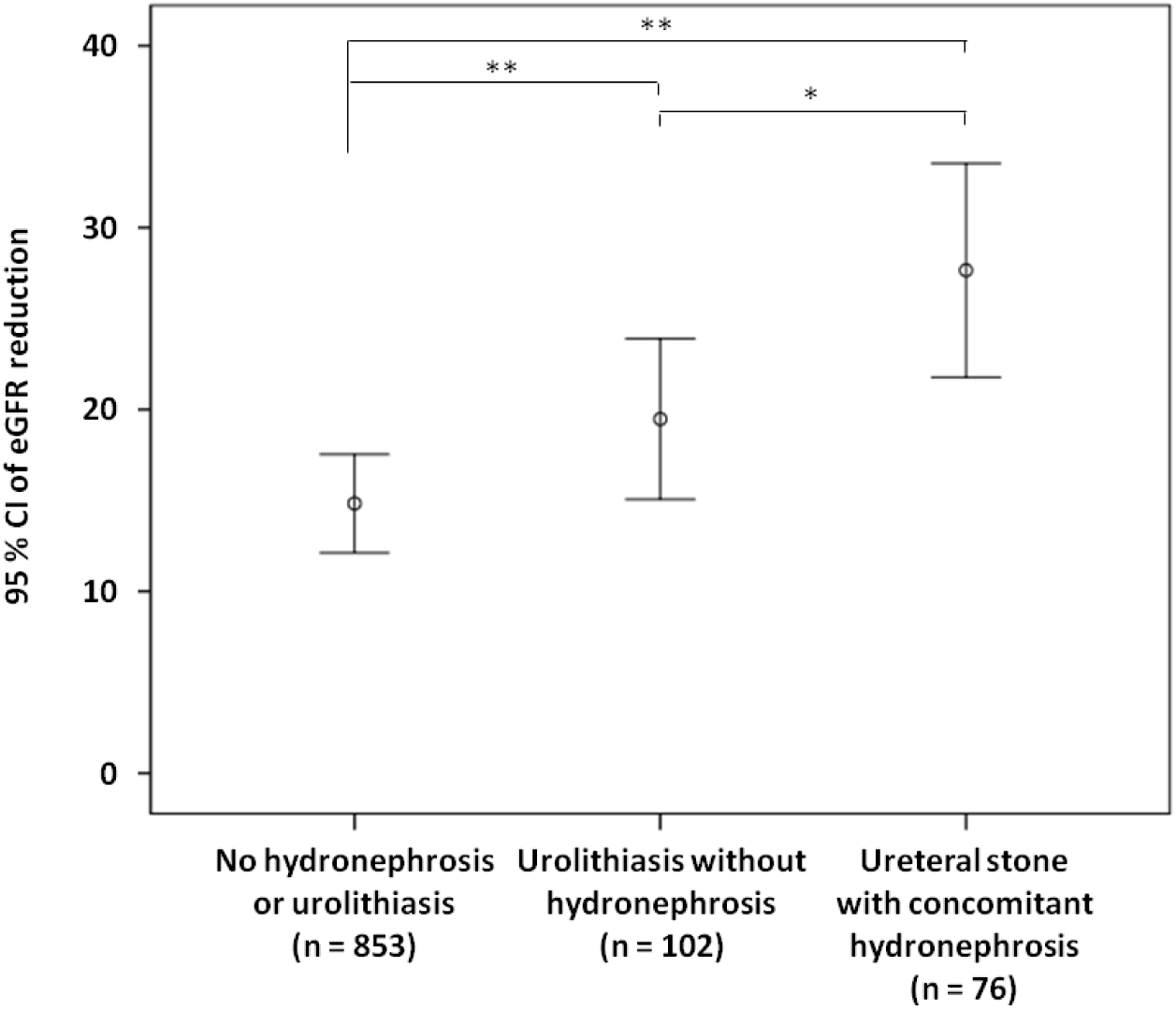
eGFR reduction in UTI patients with respect to ureteral stone with concomitant hydronephrosis and urolithiasis without hydronephrosis. eGFR: estimated glomerular filtration rate. UTI: urinary tract infection. eGFR reduction: baseline minus worst value of eGFR. *P <0.05, **P <0.01.

### Risk factors of hospitalized UTI patients with urolithiasis with respect to ureteral stone and concomitant hydronephrosis

The demographic and clinical characteristics of UTI patients with urolithiasis are shown in Table 2. Patients with ureteral stone and concomitant hydronephrosis had higher serum creatinine level on admission (2.35 ± 2.28 versus 1.58 ± 1.41 mg/mL, P = 0.02), longer hospital stay (12 ± 6 versus 9 ± 4 days, P < 0.001), higher prevalence of bacteremia (69.7% versus 49.0%, P = 0.006) and AKI (40.8% versus 22.5%, P = 0.009), and lower prevalence of afebrile status (19.7% versus 40.2%, P = 0.004) than those without ureteral stone and concomitant hydronephrosis. Univariate logistic regression analysis showed that older age (odds ratio [OR] 1.032, 95% CI 1.006–1.057, P = 0.014), uroseptic shock (OR 6.249, 95% CI 3.074–12.702, P < 0.001), and ureteral stone with concomitant hydronephrosis (OR 2.366, 95% CI 1.233–4.541, P = 0.01) were associated with an increased risk for AKI, whereas a higher platelet count (OR 0.995, 95% CI 0.991–1.000, P = 0.042) was associated with a decreased risk for AKI in UTI patients with urolithiasis. Multivariate logistic regression analysis revealed that older age (OR 1.032, 95% CI 1.002–1.063, P = 0.038), uroseptic shock (OR 4.769, 95% CI 2.184–10.412, P < 0.001), and ureteral stone with concomitant hydronephrosis (OR 2.345, 95% CI 1.128–4.876, P =0.023) were independently associated with an increased risk for AKI in UTI patients with urolithiasis (Table 3).

**Table 2.** Characteristics of hospitalized urinary tract infection patients with urolithiasis with respect to ureteral stone with hydronephrosis

| Covariate | All<br>(n = 178) | Ureteral stone with hydronephrosis |  | P-Value |
| --- | --- | --- | --- | --- |
|  |  | Non<br>(n=102) | Yes<br>(n=76) |  |
| Age (years) | 67±14 | 66±16 | 67±11 | 0.717* |
| Gender (male) | 60 (33.7) | 39 (38.2) | 21 (27.6) | 0.139<sup>¥</sup> |
| Diabetes mellitus | 81 (45.5) | 48 (47.1) | 33 (43.4) | 0.630<sup>¥</sup> |
| Hypertension | 101 (56.7) | 60 (58.8) | 41 (53.9) | 0.516<sup>¥</sup> |
| Congestive heart failure | 8 (4.5) | 5 (4.9) | 3 (3.9) | 1.000<sup>¥</sup> |
| Coronary artery disease | 20 (11.2) | 11 (10.8) | 9 (11.8) | 0.825<sup>¥</sup> |
| Stroke | 37 (20.8) | 18 (17.6) | 19 (25) | 0.232<sup>¥</sup> |
| Prior history of UTI |  |  |  | 0.638<sup>¥</sup> |
| None | 104 (58.4) | 56 (54.9) | 48 (63.2) |  |
| Once | 38 (21.3) | 25 (24.5) | 13 (17.1) |  |
| Twice | 16 (9.0) | 9 (8.8) | 7 (9.2) |  |
| Thrice or more | 20 (11.2) | 12 (11.8) | 8 (10.5) |  |
| Indwelling Foley catheter | 13 (7.3) | 8 (7.8) | 5 (6.6) | 0.748<sup>¥</sup> |
| Afebrile | 56 (31.5) | 41 (40.2) | 15 (19.7) | 0.004<sup>¥</sup> |
| Bacteremia | 103 (57.9) | 50 (49.0) | 53 (69.7) | 0.006<sup>¥</sup> |
| Uroseptic shock | 53 (29.8) | 26 (25.5) | 27 (35.5) | 0.148<sup>¥</sup> |
| Acute kidney injury | 54 (30.3) | 23 (22.5) | 31 (40.8) | 0.009<sup>¥</sup> |
| Length of hospital stay (days) | 11±5 | 9±4 | 12±6 | <0.001<sup>¥</sup> |
| All-cause in-hospital mortality | 1 (0.6) | 0 (0.0) | 1 (1.3) | 0.427<sup>¥</sup> |
| Hospitalized serum creatinine (mg/dL) | 1.91±1.86 | 1.58±1.41 | 2.35±2.28 | 0.020* |
| Baseline eGFR (mL/min/1.73 m<sup>2</sup>) | 74.24±27.59 | 76.36±26.61 | 71.40±28.78 | 0.237* |
| Mean white blood cell (10<sup>3</sup>/μL) | 13.94±7.14 | 13.82±6.24 | 14.09±8.23 | 0.803* |
| Platelets (10<sup>3</sup>/μL) | 196.31±80.96 | 200.12±84.17 | 191.21±76.70 | 0.469* |
| *Escherichia coli* | 124 (69.7) | 74 (72.5) | 50 (65.8) | 0.332<sup>¥</sup> |
| *Proteus* species | 17 (9.6) | 8 (7.8) | 9 (11.8) | 0.369<sup>¥</sup> |
| *Klebsiella* species | 14 (7.9) | 7 (6.9) | 7 (9.2) | 0.565<sup>¥</sup> |
| *Enterococcus* species | 6 (3.4) | 2 (2.0) | 4 (5.3) | 0.404<sup>¥</sup> |
| *Pseudomonas* species | 8 (4.5) | 5 (4.9) | 3 (3.9) | 1.000<sup>¥</sup> |
| Multidrug-resistance bacteria isolate | 65 (36.5) | 37 (36.3) | 28 (36.8) | 0.938<sup>¥</sup> |

Data are expressed as mean ± SD or number (percentage) \*: Student T-test or Mann-Whitney U-test; <sup>¥</sup>: Chi-Square test or Fisher's exact test
UTI: urinary tract infection; eGFR: estimated glomerular filtration rate.

**Table 3.** Univariate and multivariate logistic regression analyses of factors related to acute kidney injury in hospitalized urinary tract infection patients with urolithiasis

| Covariate | Univariate |  |  | Multivariate |  |  |
| --- | --- | --- | --- | --- | --- | --- |
| | $\beta$ | OR(95% C.I) | P-Value | $\beta$ | OR(95% C.I) | P-Value |
| Age (years) | 0.031 | 1.032 (1.006-1.057) | 0.014 | 0.031 | 1.032 (1.002-1.063) | 0.038 |
| Gender (male) | -0.144 | 0.865 (0.437-1.714) | 0.679 |  |  |  |
| Diabetes mellitus | 0.046 | 1.047 (0.551-1.988) | 0.889 |  |  |  |
| Hypertension | 0.148 | 1.159 (0.606-2.217) | 0.655 |  |  |  |
| Congestive heart failure | 0.875 | 2.400 (0.577-9.975) | 0.228 |  |  |  |
| Coronary artery disease | -0.018 | 0.982 (0.356-2.709) | 0.972 |  |  |  |
| Stroke | 0.430 | 1.537 (0.720-3.282) | 0.267 |  |  |  |
| Indwelling Foley catheter | 0.022 | 1.022 (0.301-3.475) | 0.972 |  |  |  |
| Afebrile | 0.363 | 1.438 (0.732-2.824) | 0.292 |  |  |  |
| Bacteremia | 0.417 | 1.517 (0.783-2.938) | 0.217 |  |  |  |
| Uroseptic shock | 1.832 | 6.249 (3.074-12.702) | <0.001 | 1.562 | 4.769 (2.184-10.412) | <0.001 |
| Ureteral stone with hydronephrosis | 0.861 | 2.366 (1.233-4.541) | 0.010 | 0.852 | 2.345 (1.128-4.876) | 0.023 |
| Baseline eGFR (mL/min/1.73 m<sup>2</sup>) | -0.001 | 0.999 (0.988-1.011) | 0.877 |  |  |  |
| Mean white blood cell (10<sup>3</sup>/μL) | 0.043 | 1.044 (0.998-1.092) | 0.062 | 0.032 | 1.032 (0.975-1.093) | 0.277 |
| Platelets (10<sup>3</sup>/μL) | -0.005 | 0.995 (0.991-1.000) | 0.042 | -0.004 | 0.996 (0.991-1.001) | 0.095 |
| Multidrug-resistance bacteria isolate | 0.482 | 1.620 (0.842-3.116) | 0.149 | 0.178 | 1.195 (0.559-2.554) | 0.646 |

Multivariate model: Cox & Snell $R^2 = 0.203$ Nagelkerke $R^2 = 0.287$

### Risk factors of hospitalized UTI patients without hydronephrosis with respect to urolithiasis

The demographic and clinical characteristics of UTI patients without hydronephrosis are shown in Table 4. Patients with urolithiasis had a higher prevalence of male gender (38.2% versus 25.3%, P = 0.005), prior history of UTI (45.1% versus 31.4%, P = 0.016), AKI (22.5% versus 9.6%, P < 0.001), and *Proteus* species (7.8% versus 1.6%, P = 0.001), but a lower prevalence of *E. coli* (72.5% versus 80.9%, P = 0.047), than those without urolithiasis. Univariate logistic regression analysis showed that older age (OR 1.029, 95% CI 1.015–1.043, P < 0.001), DM (OR 1.549, 95% CI 1.031–2.326, P = 0.035), hypertension (OR 2.596, 95% CI 1.665–4.048, P < 0.001), CHF (OR 2.691, 95% CI 1.282–5.646, P = 0.009), afebrile status (OR 1.556, 95% CI 1.034–2.340, P = 0.034), bacteremia (OR 1.696, 95% CI 1.128–2.550, P = 0.011), uroseptic shock (OR 4.318, 95% CI 2.814–6.627, P < 0.001), urolithiasis (OR 2.737, 95% CI 1.632–4.591, P < 0.001), and higher WBC count (OR 1.059, 95% CI 1.027–1.092, P < 0.001) were associated with an increased risk for AKI, whereas a higher baseline eGFR (OR 0.982, 95% CI 0.976–0.989, P < 0.001) was associated with a decreased risk for AKI in UTI patients without hydronephrosis. Multivariate logistic regression analysis revealed that hypertension (OR 2.064, 95% CI 1.226–3.475, P = 0.006), uroseptic shock (OR 4.505, 95% CI 2.798–7.253, P < 0.001), urolithiasis (OR 2.484, 95% CI 1.398–4.415, P = 0.002), and higher WBC count (OR 1.061, 95% CI 1.027–1.096, P < 0.001) were independently associated with an increased risk for AKI in UTI patients without hydronephrosis. Conversely, a higher baseline eGFR (OR 0.985, 95% CI 0.977–0.994, P = 0.001) was independently associated with a decreased risk for AKI in UTI patients without hydronephrosis (Table 5).

**Table 4.** Characteristics of hospitalized urinary tract infection patients without hydronephrosis with respect to urolithiasis

|  | All<br>(n = 955) | Non-Urolithiasis<br>(n=853) | Urolithiasis<br>(n=102) | P-Value |
| --- | --- | --- | --- | --- |
| Age (years) | 66±18 | 66±18 | 66±16 | 0.875* |
| Gender (male) | 255 (26.7) | 216 (25.3) | 39 (38.2) | 0.005¥ |
| Diabetes mellitus | 408 (42.7) | 360 (42.2) | 48 (47.1) | 0.349¥ |
| Hypertension | 492 (51.5) | 432 (50.6) | 60 (58.8) | 0.118¥ |
| Congestive heart failure | 42 (4.4) | 37 (4.3) | 5 (4.9) | 0.797¥ |
| Coronary artery disease | 105 (11.0) | 94 (11.0) | 11 (10.8) | 0.943¥ |
| Stroke | 205 (21.5) | 187 (21.9) | 18 (17.6) | 0.320¥ |
| Prior history of UTI |  |  |  | 0.016¥ |
| None | 641 (67.1) | 585 (68.6) | 56 (54.9) |  |
| Once | 180 (18.8) | 155 (18.2) | 25 (24.5) |  |
| Twice | 75 (7.9) | 66 (7.7) | 9 (8.8) |  |
| Thrice or more | 59 (6.2) | 47 (5.5) | 12 (11.8) |  |
| Indwelling Foley catheter | 62 (6.5) | 54 (6.3) | 8 (7.8) | 0.558¥ |
| Afebrile | 355 (37.2) | 314 (36.8) | 41 (40.2) | 0.504¥ |
| Bacteremia | 416 (43.6) | 366 (42.9) | 50 (49.0) | 0.239¥ |
| Uroseptic shock | 176 (18.4) | 150 (17.6) | 26 (25.5) | 0.052¥ |
| Acute kidney injury | 105 (11.0) | 82 (9.6) | 23 (22.5) | <0.001¥ |
| Length of hospital stay (days) | 9±5 | 9±5 | 9±4 | 0.861* |
| All-cause in-hospital mortality | 6 (0.6) | 6 (0.7) | 0 (0.0) | 0.507¥ |
| Hospitalized serum creatinine (mg/dL) | 1.47±1.39 | 1.46±1.39 | 1.58±1.41 | 0.392* |
| Baseline eGFR (mL/min/1.73 m<sup>2</sup>) | 76.84±30.14 | 76.89±30.55 | 76.36±26.61 | 0.865* |
| Mean white blood cell (10<sup>3</sup>/μL) | 13.28±5.94 | 13.21±5.91 | 13.82±6.24 | 0.330* |
| Platelets (10<sup>3</sup>/μL) | 205.56±112.65 | 206.21±115.61 | 200.12±84.17 | 0.606* |
| *Escherichia coli* | 764 (80.0) | 690 (80.9) | 74 (72.5) | 0.047¥ |
| *Proteus* species | 22 (2.3) | 14 (1.6) | 8 (7.8) | 0.001¥ |
| *Klebsiella* species | 67 (7.0) | 60 (7.0) | 7 (6.9) | 0.949¥ |
| *Enterococcus* species | 32 (3.4) | 30 (3.5) | 2 (2.0) | 0.567¥ |
| *Pseudomonas* species | 48 (5.0) | 43 (5.0) | 5 (4.9) | 0.952¥ |
| Multidrug-resistance bacteria isolate | 327 (34.2) | 290 (34.0) | 37 (36.3) | 0.647¥ |

Data are expressed as mean ± SD or number (percentage) \*: Student T-test; ¥:Chi-Square test or Fisher's exact test UTI: urinary tract infection; eGFR: estimated glomerular filtration rate.

**Table 5.** Univariate and multivariate logistic regression analyses of factors related to acute kidney injury in urinary tract infection patients without hydronephrosis

| Covariate | Univariate |  |  | Multivariate |  |  |
| --- | --- | --- | --- | --- | --- | --- |
| | $\beta$ | OR(95% C.I) | P-Value | $\beta$ | OR(95% C.I) | P-Value |
| Age (years) | 0.029 | 1.029 (1.015-1.043) | <0.001 | 0.008 | 1.008 (0.990-1.027) | 0.362 |
| Gender (male) | 0.260 | 1.297 (0.835-2.012) | 0.247 |  |  |  |
| Diabetes mellitus | 0.437 | 1.549 (1.031-2.326) | 0.035 | 0.122 | 1.130 (0.713-1.791) | 0.604 |
| Hypertension | 0.954 | 2.596 (1.665-4.048) | <0.001 | 0.725 | 2.064 (1.226-3.475) | 0.006 |
| Congestive heart failure | 0.990 | 2.691 (1.282-5.646) | 0.009 | 0.399 | 1.491 (0.629-3.535) | 0.365 |
| Coronary artery disease | 0.514 | 1.673 (0.952-2.941) | 0.074 | -0.067 | 0.936 (0.487-1.796) | 0.841 |
| Stroke | 0.210 | 1.234 (0.769-1.980) | 0.384 |  |  |  |
| Indwelling Foley catheter | 0.480 | 1.615 (0.795-3.284) | 0.185 |  |  |  |
| Afebrile | 0.442 | 1.556 (1.034-2.340) | 0.034 | 0.422 | 1.525 (0.948-2.455) | 0.082 |
| Bacteremia | 0.528 | 1.696 (1.128-2.550) | 0.011 | 0.465 | 1.591 (0.994-2.548) | 0.053 |
| Uroseptic shock | 1.463 | 4.318 (2.814-6.627) | <0.001 | 1.505 | 4.505 (2.798-7.253) | <0.001 |
| Urolithiasis | 1.007 | 2.737 (1.632-4.591) | <0.001 | 0.910 | 2.484 (1.398-4.415) | 0.002 |
| Baseline eGFR (mL/min/1.73 m<sup>2</sup>) | -0.018 | 0.982 (0.976-0.989) | <0.001 | -0.015 | 0.985 (0.977-0.994) | 0.001 |
| Mean white blood cell (10<sup>3</sup>/μL) | 0.058 | 1.059 (1.027-1.092) | <0.001 | 0.059 | 1.061 (1.027-1.096) | <0.001 |
| Platelets (10<sup>3</sup>/μL) | -0.002 | 0.998 (0.996-1.001) | 0.151 |  |  |  |
| Multidrug-resistance bacteria isolate | 0.279 | 1.322 (0.872-2.003) | 0.188 |  |  |  |

Multivariate model: Cox & Snell $R^2 = 0.112$ Nagelkerke $R^2 = 0.223$

## Discussion

UTI is a common complication in patients with urolithiasis. Urolithiasis is a common cause of urinary tract obstruction predisposing to UTI [15]. A previous study showed that urinary obstruction is a predictor of the development of septic shock in UTI patients [16]. In this study, we identified that the presence of ureteral stone with hydronephrosis is a risk factor for AKI in UTI patients with urolithiasis. In addition, urolithiasis is a risk factor for AKI in UTI patients without hydronephrosis. To our knowledge, this is the first study to demonstrate that urolithiasis without hydronephrosis is associated with an increased risk for AKI in patients with UTI.

Urolithiasis is a major cause of obstructive pyelonephritis, accounting for 65% of obstructive pyelonephritis events [17]. Urinary stasis provides the time and opportunity for bacteria to adhere to the urothelium, multiply, and infect the host [18]. In addition, stones moving down the ureter result in inflamed narrowing of the ureter or injuries, which could easily cause infection [19]. Obstructive uropathy is not a rare cause of UTI. A previous study showed that the rate of obstructive uropathy in patients clinically diagnosed with a simple pyelonephritis was approximately 6% [20]. Obstructive UTI is associated with a high recurrence rate. In a long-term follow-up study in patients with obstructive pyelonephritis, the incidence of recurrent obstructive pyelonephritis and recurrent UTI was 11% and 33%, respectively [17]. The existence of ureteral stone is the most common cause of urosepsis in patients with urinary tract obstruction [16]. UTI in the presence of urinary tract obstruction is a serious clinical situation. In the case of acute obstructive uropathy, the increased intrarenal pelvic pressure decreases the drug delivery to the kidney [21]. Both urolithiasis and obstructive uropathy have an impact on the severity of urosepsis [22], the presence of urinary tract obstruction in patients with acute pyelonephritis is associated with an increased risk for urosepsis and uroseptic shock [23, 24]. In the current study, the presence of ureteral stone with hydronephrosis was an independent risk factor for AKI development in UTI patients with urolithiasis.

Obstructive nephropathy is the major cause of non-infectious urolithiasis associated AKI [25]. Renal vasoconstriction and inflammation can occur in response to increased intratubular pressure, and cause ischemia injury. If ischemia persists, glomerulosclerosis, tubular atrophy, and interstitial fibrosis occur [26]. In a rat model, complete unilateral obstruction for 24 hours resulted in irreversible loss of function in 15% of the nephrons of the affected kidney [27]. Obstructive AKI caused by urolithiasis is not common, accounting for only approximately 1–2% of all AKI events [25]. A previous study reported that the prevalence of AKI among patients with non-infectious ureteral stone was 0.72% [28]. Infection is one of the leading causes of AKI in patients with ureteral stone [29], and infection with concomitant obstructive uropathy causes loss of function far more rapidly than a simple obstructive uropathy [30]. Yamamoto et al. found that approximately 80% of patients with acute obstructive pyelonephritis induced by upper ureteral stone had elevated serum creatinine levels above the normal range [31]. AKI is associated with an increased risk for mortality in patients with upper ureteral stone with concomitant urosepsis [32]. Untreated infection with concomitant urinary tract obstruction can lead to life-threatening sepsis. After the administration of appropriate antibiotics, interventions, such as drainage for source control, should be performed within 12 hours after the diagnosis of sepsis, according to the recommendation from the Surviving Sepsis Campaign 2016 [33]. The management of infected hydronephrosis secondary to urolithiasis is prompt decompression of the renal collecting system [34]. Optional methods of decompression include percutaneous nephrostomy and retrograde ureteral stenting [22]. Decompression of the renal collecting system increases the renal plasma flow and delivery of antibiotics to both the renal parenchyma and urine [35]. Moreover, the release of inner pressure from the upper urinary tracts reduces the invasion of bacteria into the bloodstream or renal parenchyma [36]. Both the increase of antibiotic penetration into the affected renal unit and the decrease of bacterial invasion can facilitate infection control. Further, the severity of tubular atrophy and interstitial fibrosis increases if the duration of obstruction is extended [37], and a delayed relief of ureteral obstruction decreases long-term renal function [38]. If obstruction is relieved early and infection is controlled, the renal parenchyma has a great potential for recovery [29]. Delay or omission of renal decompression will increase the rate of sepsis and mortality [17]. Because UTI patients with concomitant ureteral stone resulting in urinary tract obstruction are predisposed to develop severe complications, such as septic shock or AKI, timely decompression of the renal collecting system through percutaneous nephrostomy or retrograde ureteral stenting after starting empiric antibiotic therapy is recommended for those with a critical status [39].

Urolithiasis is a source of infection [40], and UTIs are frequently associated with almost all chemical types of kidney stones [41]. Urolithiasis is the one of most common urological disorders associated with complicated pyelonephritis [42] and is also a risk factor for treatment failure in patients with acute pyelonephritis [43]. Because the amount of bacteria on the stone surface can increase despite antibiotic therapy, eradication of the associated UTI is only possible after the stone has been completely removed [44]. According to the Nationwide Inpatient Sample survey between 1999 and 2009, the incidence of urolithiasis-associated sepsis and severe sepsis is increasing [45]. Urolithiasis is also a risk factor of septic shock in patients with UTI [7], and sepsis is one of the most common etiologies of AKI [46]. In the current study, urolithiasis without hydronephrosis was associated with an increased risk for AKI in patients with UTI. Early imaging examination to identify urolithiasis and complete removal of urolithiasis after ameliorating the symptoms of sepsis are crucial for patients with UTI.

Our study had several limitations. First, this was a single-center study with a retrospective design. A multicenter prospective study with a larger sample size will be needed to confirm our results. Second, we did not routinely examine the stone characteristics in this study. Further studies evaluating the chemical types of stones will be needed to confirm their impact on AKI in patients with UTI.

## Conclusions

We demonstrated that patients with ureteral stone complicated with hydronephrosis have a higher risk of developing AKI among UTI patients with urolithiasis, and urolithiasis remains a risk factor for AKI among UTI patients without hydronephrosis. Therefore, timely decompression of the renal collecting system for infectious hydronephrosis secondary to urolithiasis and complete stone removal after controlling infection are crucial for UTI patients with urolithiasis.

## Acknowledgments

The authors would like to acknowledge the support of Cheng-Lun Chiang.

## Author Contributions

**Conceptualization:** Chih-Yen Hsiao.

**Data curation:** Chih-Yen Hsiao.

**Formal analysis:** Yi-Chien Lee and Tsung-Hsien Chen.

**Methodology:** Chih-Yen Hsiao.

**Supervision:** Ming-Cheng Wang.

**Writing – original draft:** Chih-Yen Hsiao, and Tsung-Hsien Chen.

**Writing – review & editing:** Ming-Cheng Wang.

## References

1. Coe FL, Evan A, Worcester E. Kidney stone disease. J Clin Invest. 2005;115(10):2598–608. doi: 10.1172/JCI26662. PubMed PMID: 16200192; PubMed Central PMCID: PMCPMC1236703.

2. Worcester EM, Coe FL. Clinical practice. Calcium kidney stones. N Engl J Med. 2010;363(10):954–63. doi: 10.1056/NEJMcp1001011. PubMed PMID: 20818905; PubMed Central PMCID: PMCPMC3192488.

3. Scales CD, Jr., Smith AC, Hanley JM, Saigal CS, Urologic Diseases in America P. Prevalence of kidney stones in the United States. Eur Urol. 2012;62(1):160–5. doi: 10.1016/j.eururo.2012.03.052. PubMed PMID: 22498635; PubMed Central PMCID: PMCPMC3362665.

4. Sakhaee K. Nephrolithiasis as a systemic disorder. Curr Opin Nephrol Hypertens. 2008;17(3):304–9. doi: 10.1097/MNH.0b013e3282f8b34d. PubMed PMID: 18408483.

5. Moe OW. Kidney stones: pathophysiology and medical management. Lancet. 2006;367(9507):333–44. doi: 10.1016/S0140-6736(06)68071-9. PubMed PMID: 16443041.

6. Baker LR, Cattell WR, Fry IK, Mallinson WJ. Acute renal failure due to bacterial pyelonephritis. Q J Med. 1979;48(192):603-12. PubMed PMID: 395560.

7. Hsiao CY, Chen TH, Lee YC, Hsiao MC, Hung PH, Chen YY, et al. Urolithiasis Is a Risk Factor for Uroseptic Shock and Acute Kidney Injury in Patients With Urinary Tract Infection. Front Med. 2019;6:288. doi: 10.3389/fmed.2019.00288. PubMed PMID: 31867338; PubMed Central PMCID: PMCPMC6906152.

8. Liborio AB, Leite TT, Neves FM, Teles F, Bezerra CT. AKI complications in critically ill patients: association with mortality rates and RRT. Clin J Am Soc Nephrol. 2015;10(1):21–8. doi: 10.2215/CJN.04750514. PubMed PMID: 25376761; PubMed Central PMCID: PMCPMC4284413.

9. Levey AS, Stevens LA, Schmid CH, Zhang YL, Castro AF, 3rd, Feldman HI, et al. A new equation to estimate glomerular filtration rate. Ann Intern Med. 2009;150(9):604-12. PubMed PMID: 19414839; PubMed Central PMCID: PMCPMC2763564.

10. National Kidney F. K/DOQI clinical practice guidelines for chronic kidney disease: evaluation, classification, and stratification. Am J Kidney Dis. 2002;39(2 Suppl 1):S1-266. PubMed PMID: 11904577.

11. Kellum J, Lameire N, Aspelin P, Barsoum R, Burdmann E, Goldstein S, et al. Kidney disease: Improving global outcomes (KDIGO) acute kidney injury work group. KDIGO clinical practice guideline for acute kidney injury. Kidney Int Suppl. 2012;2:1–138.

12. Dellinger RP, Levy MM, Rhodes A, Annane D, Gerlach H, Opal SM, et al. Surviving sepsis campaign: international guidelines for management of severe sepsis and septic shock: 2012. Crit Care Med. 2013;41(2):580–637. doi: 10.1097/CCM.0b013e31827e83af. PubMed PMID: 23353941.

13. Dalal S, Zhukovsky DS. Pathophysiology and management of fever. J Support Oncol. 2006;4(1):9-16. PubMed PMID: 16444847.

14. Magiorakos AP, Srinivasan A, Carey RB, Carmeli Y, Falagas ME, Giske CG, et al. Multidrug-resistant, extensively drug-resistant and pandrug-resistant bacteria: an international expert proposal for interim standard definitions for acquired resistance. Clin Microbiol Infect. 2012;18(3):268–81. doi: 10.1111/j.1469-0691.2011.03570.x. PubMed PMID: 21793988.

15. Cohen J, Cohen S, Grasso M. Ureteropyeloscopic treatment of large, complex intrarenal and proximal ureteral calculi. BJU Int. 2013;111(3 Pt B):E127-31. doi: 10.1111/j.1464-410X.2012.11352.x. PubMed PMID: 22757752.

16. Wagenlehner FME, Pilatz A, Weidner W, Naber KG. Urosepsis: Overview of the Diagnostic and Treatment Challenges. Microbiol Spectr. 2015;3(5). doi: 10.1128/microbiolspec.UTI-0003-2012. PubMed PMID: 26542042.

17. Vahlensieck W, Friess D, Fabry W, Waidelich R, Bschleipfer T. Long-term results after acute therapy of obstructive pyelonephritis. Urol Int. 2015;94(4):436–41. doi: 10.1159/000368051. PubMed PMID: 25661913.

18. Cox CE, Hinman F, Jr. Experiments with induced bacteriuria, vesical emptying and bacterial growth on the mechanism of bladder defense to infection. J Urol. 1961;86:739–48. doi: 10.1016/s0022-5347(17)65257-1. PubMed PMID: 13881887.

19. Yongzhi L, Shi Y, Jia L, Yili L, Xingwang Z, Xue G. Risk factors for urinary tract infection in patients with urolithiasis-primary report of a single center cohort. BMC Urol. 2018;18(1):45. doi: 10.1186/s12894-018-0359-y. PubMed PMID: 29783970; PubMed Central PMCID: PMCPMC5963162.

20. Lujan Galan M, Paez Borda A, Fernandez Gonzalez I, Llorente Abarca C, Romero Cajigal I, Bustamante Alarma S, et al. [Usefulness of ultrasonography in the assessment of acute pyelonephritis]. Arch Esp Urol. 1997;50(1):46-50. PubMed PMID: 9182488.

21. Berger I, Wildhofen S, Lee A, Ponholzer A, Rauchenwald M, Zechner O, et al. Emergency nephrectomy due to severe urosepsis: a retrospective, multicentre analysis of 65 cases. BJU Int. 2009;104(3):386–90. doi: 10.1111/j.1464-410X.2009.08414.x. PubMed PMID: 19338556.

22. Tambo M, Okegawa T, Shishido T, Higashihara E, Nutahara K. Predictors of septic shock in obstructive acute pyelonephritis. World J Urol. 2014;32(3):803–11. doi: 10.1007/s00345-013-1166-4. PubMed PMID: 24037335; PubMed Central PMCID: PMCPMC4031390.

23. Lee JH, Lee YM, Cho JH. Risk factors of septic shock in bacteremic acute pyelonephritis patients admitted to an ER. tJ Infect Chemother. 2012;18(1):130–3. doi: 10.1007/s10156-011-0289-z. PubMed PMID: 21861118.

24. Yamamichi F, Shigemura K, Kitagawa K, Fujisawa M. Comparison between non-septic and septic cases in stone-related obstructive acute pyelonephritis and risk factors for septic shock: A multi-center retrospective study. J Infect Chemother. 2018;24(11):902–6. doi: 10.1016/j.jiac.2018.08.002. PubMed PMID: 30174285.

25. Tang X, Lieske JC. Acute and chronic kidney injury in nephrolithiasis. Curr Opin Nephrol Hypertens. 2014;23(4):385–90. doi: 10.1097/01.mnh.0000447017.28852.52. PubMed PMID: 24848936; PubMed Central PMCID: PMCPMC4096690.

26. Rule AD, Krambeck AE, Lieske JC. Chronic kidney disease in kidney stone formers. Clin J Am Soc Nephrol. 2011;6(8):2069–75. doi: 10.2215/CJN.10651110. PubMed PMID: 21784825; PubMed Central PMCID: PMCPMC3156433.

27. Bander SJ, Buerkert JE, Martin D, Klahr S. Long-term effects of 24-hr unilateral ureteral obstruction on renal function in the rat. Kidney Int. 1985;28(4):614–20. doi: 10.1038/ki.1985.173. PubMed PMID: 4087683.

28. Wang SJ, Mu XN, Zhang LY, Liu QY, Jin XB. The incidence and clinical features of acute kidney injury secondary to ureteral calculi. Urol Res. 2012;40(4):345–8. doi: 10.1007/s00240-011-0414-6. PubMed PMID: 21853241.

29. Singh SM, Yadav R, Gupta NP, Wadhwa SN. The management of renal and ureteric calculi in renal failure. Br J Urol. 1982;54(5):455–7. doi: 10.1111/j.1464-410x.1982.tb13563.x. PubMed PMID: 7171948.

30. Holm-Nielsen A, Jorgensen T, Mogensen P, Fogh J. The prognostic value of probe renography in ureteric stone obstruction. Br J Urol. 1981;53(6):504–7. doi: 10.1111/j.1464-410x.1981.tb03248.x. PubMed PMID: 7317731.

31. Yamamoto Y, Fujita K, Nakazawa S, Hayashi T, Tanigawa G, Imamura R, et al. Clinical characteristics and risk factors for septic shock in patients receiving emergency drainage for acute pyelonephritis with upper urinary tract calculi. BMC Urol. 2012;12:4. doi: 10.1186/1471-2490-12-4. PubMed PMID: 22413829; PubMed Central PMCID: PMCPMC3353222.

32. Badia M, Iglesias S, Servia L, Domingo J, Gormaz P, Vilanova J, et al. [Mortality predictive factors in patients with urinary sepsis associated to upper urinary tract calculi]. Med Intensiva. 2015;39(5):290–7. doi: 10.1016/j.medin.2014.07.003. PubMed PMID: 25444058.

33. Rhodes A, Evans LE, Alhazzani W, Levy MM, Antonelli M, Ferrer R, et al. Surviving Sepsis Campaign: International Guidelines for Management of Sepsis and Septic Shock: 2016. Intensive Care Med. 2017;43(3):304–77. doi: 10.1007/s00134-017-4683-6. PubMed PMID: 28101605.

34. Brown PD. Management of urinary tract infections associated with nephrolithiasis. Curr Infect Dis Rep. 2010;12(6):450–4. doi: 10.1007/s11908-010-0141-0. PubMed PMID: 21308554.

35. Browne RF, Zwirewich C, Torreggiani WC. Imaging of urinary tract infection in the adult. Eur Radiol. 2004;14 Suppl 3:E168–83. doi: 10.1007/s00330-003-2050-1. PubMed PMID: 14749952.

36. Hamasuna R, Takahashi S, Nagae H, Kubo T, Yamamoto S, Arakawa S, et al. Obstructive pyelonephritis as a result of urolithiasis in Japan: diagnosis, treatment and prognosis. Int J Urol. 2015;22(3):294–300. doi: 10.1111/iju.12666. PubMed PMID: 25400222.

37. Chevalier RL, Thornhill BA, Chang AY. Unilateral ureteral obstruction in neonatal rats leads to renal insufficiency in adulthood. Kidney Int. 2000;58(5):1987–95. doi: 10.1111/j.1523-1755.2000.00371.x. PubMed PMID: 11044219.

38. Lucarelli G, Ditonno P, Bettocchi C, Grandaliano G, Gesualdo L, Selvaggi FP, et al. Delayed relief of ureteral obstruction is implicated in the long-term development of renal damage and arterial hypertension in patients with unilateral ureteral injury. J Urol. 2013;189(3):960–5. doi: 10.1016/j.juro.2012.08.242. PubMed PMID: 23017525.

39. Heyns CF. Urinary tract infections in obstructionof the urinary tract. In: Naber KG, Schaeffer AJ, Heyns CF, Matsumoto T, Shoskes DA, Bjerklund Johansen TE editors. Urogenital Infections. Arnhem: European Association of Urology-International Consultation on Urological Diseases; 2010. Pp. 452–80.

40. Zanetti G, Paparella S, Trinchieri A, Prezioso D, Rocco F, Naber KG. Infections and urolithiasis: current clinical evidence in prophylaxis and antibiotic therapy. Arch Ital Urol Androl. 2008;80(1):5-12. PubMed PMID: 18533618.

41. Tavichakorntrakool R, Prasongwattana V, Sungkeeree S, Saisud P, Sribenjalux P, Pimratana C, et al. Extensive characterizations of bacteria isolated from catheterized urine and stone matrices in patients with nephrolithiasis. Nephrol Dial Transplant. 2012;27(11):4125–30. doi: 10.1093/ndt/gfs057. PubMed PMID: 22461670.

42. Kakinoki H, Tobu S, Kakinoki Y, Udo K, Uozumi J, Noguchi M. Risk Factors for Uroseptic Shock in Patients with Urolithiasis-Related Acute Pyelonephritis. Urol Int. 2018;100(1):37–42. doi: 10.1159/000481801. PubMed PMID: 29065405.

43. Pertel PE, Haverstock D. Risk factors for a poor outcome after therapy for acute pyelonephritis. BJU Int. 2006;98(1):141–7. doi: 10.1111/j.1464-410X.2006.06222.x. PubMed PMID: 16831159.

44. Miano R, Germani S, Vespasiani G. Stones and urinary tract infections. Urol Int. 2007;79 Suppl 1:32–6. doi: 10.1159/000104439. PubMed PMID: 17726350.

45. Sammon JD, Ghani KR, Karakiewicz PI, Bhojani N, Ravi P, Sun M, et al. Temporal trends, practice patterns, and treatment outcomes for infected upper urinary tract stones in the United States. Eur Urol. 2013;64(1):85–92. doi: 10.1016/j.eururo.2012.09.035. PubMed PMID: 23031677.

46. Mori T, Shimizu T, Tani T. Septic acute renal failure. Contrib Nephrol. 2010;166:40–6. doi: 10.1159/000314849. PubMed PMID: 20472990.

